## Supplementary material for "LMSM: a modular approach for identifying lncRNA related miRNA sponge modules in breast cancer": S1 File

In the supplementary file, we will show the robustness of the LMSM workflow. In the LMSM workflow, the SGFA method [1] instead of the WGCNA method [2] is used to identify lncRNA-mRNA co-expression modules. The SGFA method is extended from the group factor analysis (GFA) method [3-5], and it can reliably infer ncRNA-mRNA co-expression modules from multiple data sources.

**Table S1. BRCA-related LMSM modules**. *L*_2_ is the number of BRCA genes in each LMSM module, *K*_2_ represents the number of genes in each LMSM module, the number of BRCA genes in the dataset (*M*_2_) is 4819, and the number of genes in the dataset (*N*_2_) is 31055. The method of identifying lncRNA-mRNA co-expression modules is the SGFA method.

| **Module ID** | ***L*_2_** | ***K*_2_** | ***p*-value** |
| --- | --- | --- | --- |
| LMSM 3 | 35 | 129 | 5.02E-04 |
| LMSM 24 | 58 | 225 | 4.67E-05 |
| LMSM 37 | 54 | 256 | 1.05E-02 |

**Table S2**. **Survival analysis of LMSM modules in BRCA**. HRlow95 and HRup95 represent the lower and upper of 95% confidence interval of HR, respectively. The identified LMSM modules can distinguish the high and the low risk BRCA samples. The method of identifying lncRNA-mRNA co-expression modules is the SGFA method.

| **Module ID** | **Chi-square** | ***p-*value** | **HR** | **HRlow95** | **HRup95** |
| --- | --- | --- | --- | --- | --- |
| LMSM 1 | 44.73 | 0 | 4.63 | 2.85 | 7.53 |
| LMSM 2 | 137.90 | 0 | 12.82 | 7.54 | 21.82 |
| LMSM 3 | 79.11 | 0 | 13.35 | 8.32 | 21.44 |
| LMSM 4 | 130.22 | 0 | 15.18 | 9.08 | 25.40 |
| LMSM 6 | 116.65 | 0 | 13.66 | 8.26 | 22.60 |
| LMSM 7 | 123.68 | 0 | 9.54 | 5.58 | 16.33 |
| LMSM 8 | 77.43 | 0 | 12.93 | 8.06 | 20.73 |
| LMSM 9 | 118.29 | 0 | 8.82 | 5.15 | 15.08 |
| LMSM 10 | 114.46 | 0 | 10.67 | 6.34 | 17.95 |
| LMSM 11 | 78.48 | 0 | 7.94 | 4.86 | 12.98 |
| LMSM 12 | 113.78 | 0 | 9.60 | 5.72 | 16.12 |
| LMSM 13 | 126.32 | 0 | 13.68 | 8.17 | 22.90 |
| LMSM 14 | 137.76 | 0 | 13.81 | 8.14 | 23.43 |
| LMSM 15 | 79.19 | 0 | 11.91 | 7.41 | 19.16 |
| LMSM 16 | 131.02 | 0 | 15.20 | 9.09 | 25.43 |
| LMSM 17 | 84.39 | 0 | 9.04 | 5.55 | 14.70 |
| LMSM 18 | 125.92 | 0 | 11.18 | 6.61 | 18.91 |
| LMSM 19 | 117.06 | 0 | 10.95 | 6.49 | 18.49 |
| LMSM 20 | 102.56 | 0 | 10.61 | 6.43 | 17.49 |
| LMSM 21 | 126.39 | 0 | 13.59 | 8.12 | 22.73 |
| LMSM 22 | 126.92 | 0 | 14.70 | 8.82 | 24.50 |
| LMSM 23 | 137.89 | 0 | 13.66 | 8.06 | 23.14 |
| LMSM 24 | 114.30 | 0 | 9.53 | 5.63 | 16.14 |
| LMSM 26 | 112.30 | 0 | 10.63 | 6.39 | 17.70 |
| LMSM 27 | 147.92 | 0 | 13.60 | 7.93 | 23.33 |
| LMSM 28 | 134.24 | 0 | 10.93 | 6.34 | 18.86 |
| LMSM 29 | 103.88 | 0 | 6.12 | 3.39 | 11.06 |
| LMSM 30 | 122.37 | 0 | 14.25 | 8.58 | 23.67 |
| LMSM 31 | 108.68 | 0 | 10.69 | 6.42 | 17.82 |
| LMSM 32 | 103.80 | 0 | 10.66 | 6.46 | 17.59 |
| LMSM 33 | 96.27 | 0 | 8.42 | 5.04 | 14.06 |
| LMSM 34 | 133.44 | 0 | 12.56 | 7.40 | 21.31 |
| LMSM 35 | 116.80 | 0 | 12.07 | 7.23 | 20.15 |
| LMSM 36 | 133.40 | 0 | 13.36 | 7.91 | 22.57 |
| LMSM 37 | 132.54 | 0 | 11.31 | 6.60 | 19.37 |
| LMSM 38 | 144.22 | 0 | 12.00 | 6.94 | 20.72 |
| LMSM 39 | 146.36 | 0 | 10.85 | 6.15 | 19.15 |
| LMSM 40 | 100.87 | 0 | 10.00 | 6.05 | 16.55 |
| LMSM 41 | 110.26 | 0 | 11.45 | 6.89 | 19.01 |
| LMSM 42 | 119.99 | 0 | 11.94 | 7.16 | 19.91 |
| LMSM 43 | 123.84 | 0 | 11.50 | 6.85 | 19.31 |
| LMSM 44 | 138.95 | 0 | 11.81 | 6.85 | 20.35 |
| LMSM 45 | 111.48 | 0 | 10.59 | 6.36 | 17.62 |
| LMSM 46 | 188.23 | 0 | 13.08 | 7.17 | 23.84 |
| LMSM 47 | 122.63 | 0 | 11.84 | 7.03 | 19.94 |
| LMSM 48 | 141.91 | 0 | 14.17 | 8.33 | 24.12 |
| LMSM 49 | 150.27 | 0 | 14.76 | 8.62 | 25.26 |
| LMSM 50 | 119.12 | 0 | 13.13 | 7.88 | 21.89 |
| LMSM 51 | 105.11 | 0 | 10.92 | 6.61 | 18.05 |

**Table S3.**  **Experimentally validated lncRNA-related miRNA sponge interactions**. The method of identifying lncRNA-mRNA co-expression modules is the SGFA method.

| **Module ID** | **Validated lncRNA- related miRNA sponge interactions** |
| --- | --- |
| LMSM 2 | *LINC00052*:*NTRK3* |
| LMSM 16 | *PVT1*:*VEGFC* |
| LMSM 27 | *SNHG7*:*CCND1* |
| LMSM 41 | *H19*:*AKT2* |

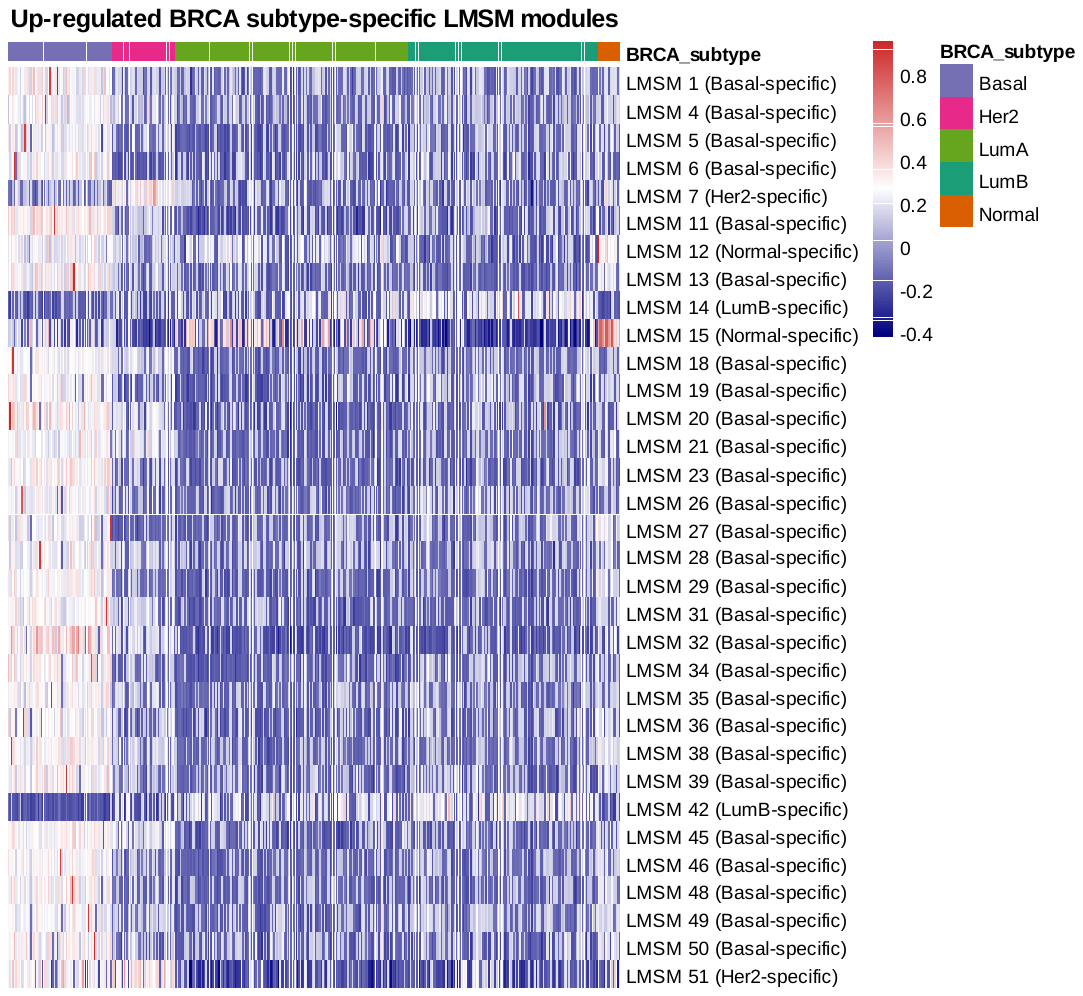

**B**

**A**

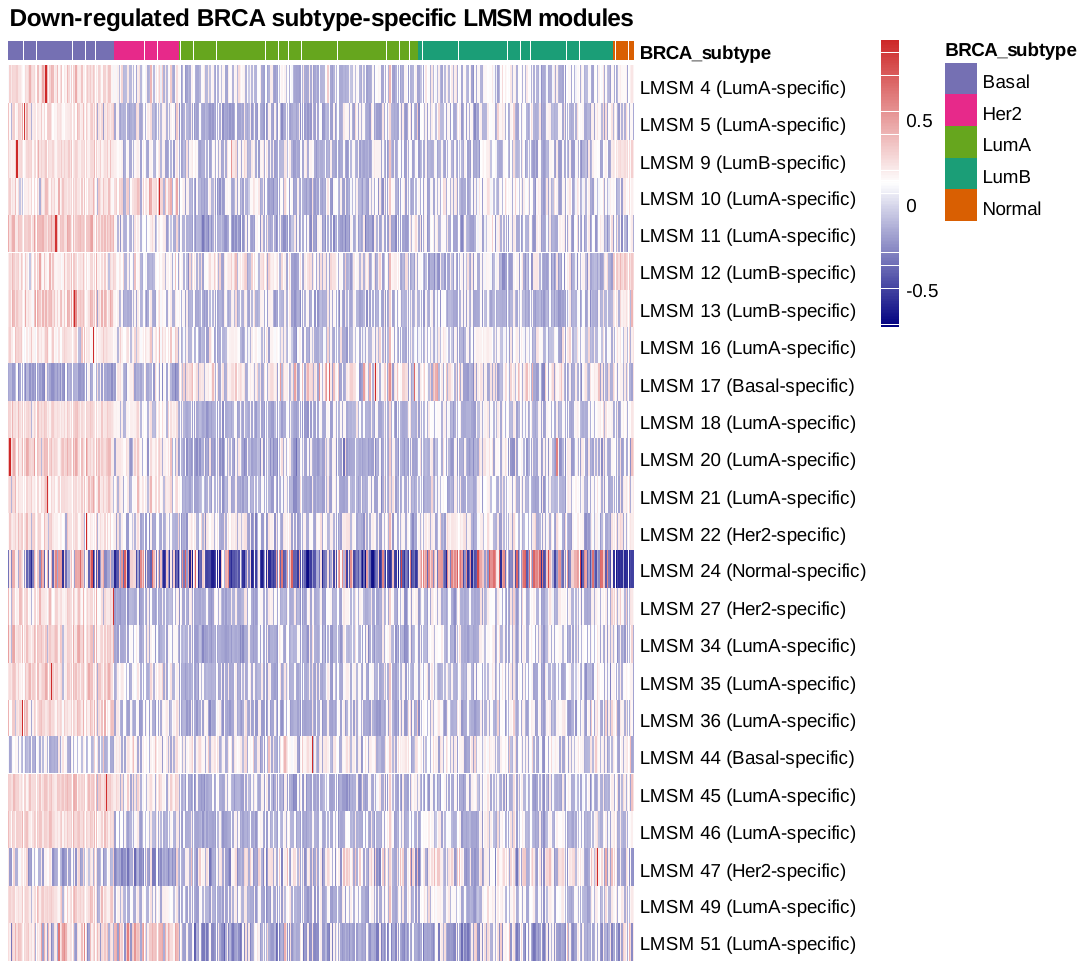

**Fig S1. Heatmap of the enrichment scores of BRCA subtype-specific LMSM modules in five BRCA subtype samples**. (A) Up-regulated BRCA subtype-specific LMSM modules. (B) Down-regulated BRCA subtype-specific LMSM modules. The method of identifying lncRNA-mRNA co-expression modules is the SGFA method.

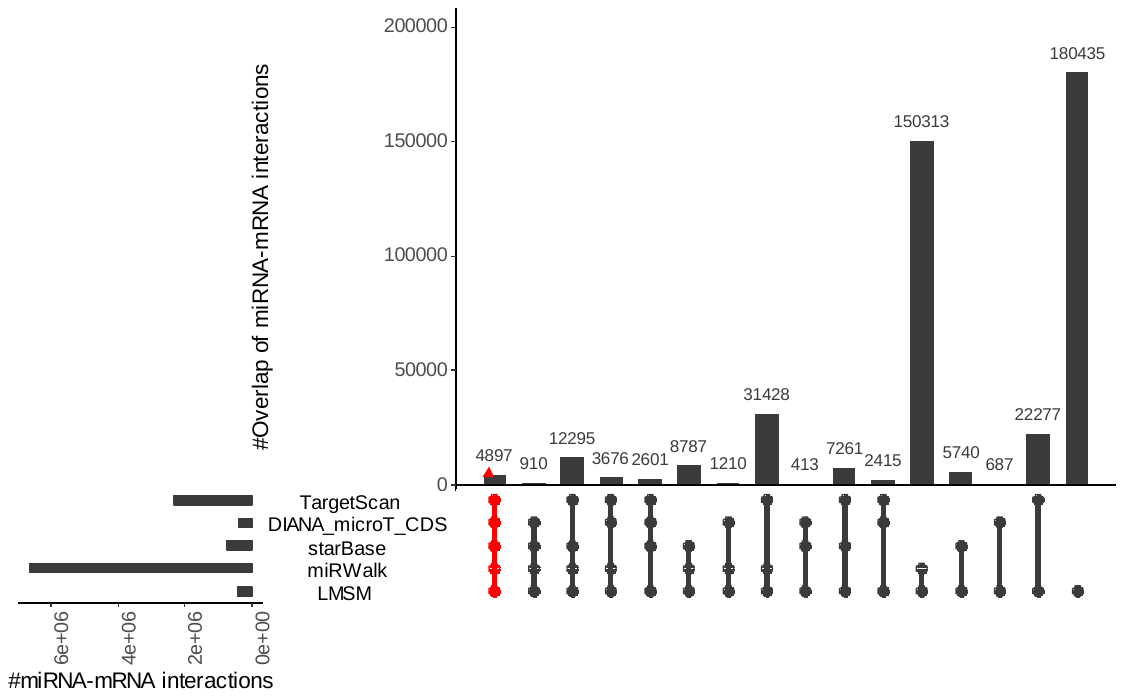

**A**

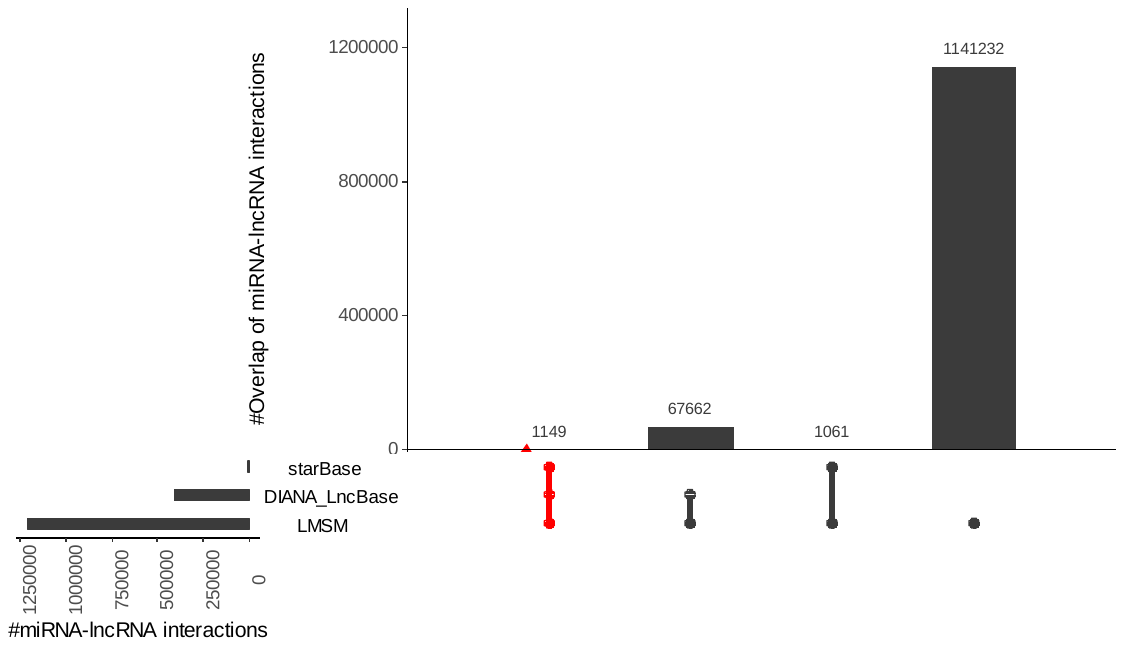

**B**

**Fig S2. Overlaps and differences between predicted miRNA-target interactions by LMSM and other methods**. (A) Predicted miRNA-mRNA interactions between LMSM and TargetScan, DIANA_microT_CDS, starBase, miRWalk. (B) Predicted miRNA-lncRNA interactions between LMSM and starBase, DIANA_LncBase. Each column corresponds to an exclusive intersection that includes the elements of the sets denoted by the dark or red circles, but not of the others. The overlap size between different methods denotes exclusive overlaps, i.e. the overlap set not in a subset of any other overlap set. The method of identifying lncRNA-mRNA co-expression modules is the SGFA method.

**Table S4. Comparison results between LMSM and GC**.

| **Method** | **%BRCA-related modules** | **%Module biomarkers** | **Mean *Subset accuracy*** | **Mean *Hamming loss*** | **#Validated interactions** |
| --- | --- | --- | --- | --- | --- |
| LMSM | 5.88% | **96.08%** | **0.6921** | **0.1135** | **4** |
| GC | **32.41%** | 66.67% | 0.6586 | 0.1319 | 2 |
